# The Dynamic Response of Human Lungs Due to Underwater Shock Wave Exposure

**DOI:** 10.1101/2023.08.08.552452

**Authors:** Eyal Bar-Kochba, Alexander S. Iwaskiw, Jenna M. Dunn, Kyle A. Ott, Timothy P. Harrigan, Constantine K. Demetropoulos

**Affiliations:** Research and Exploratory Development Department, Johns Hopkins University Applied Physics Laboratory, Laurel, MD, USA; ASRC Federal, Beltsville, MD, 20705, USA

## Abstract

Since the 19th century, underwater explosions have posed a significant threat to service members. While there have been attempts to establish injury criteria for the most vulnerable organs, namely the lungs, existing criteria are highly variable due to insufficient human data and the corresponding inability to understand the underlying injury mechanisms. This study presents an experimental characterization of isolated human lung dynamics during simulated exposure to underwater shock waves. We found that the large acoustic impedance at the surface of the lung severely attenuated transmission of the shock wave into the lungs. However, the shock wave initiated large bulk pressure-volume cycles that are distinct from the response of the solid organs under similar loading. These pressure-volume cycles are due to compression of the contained gas, which we modeled with the Rayleigh-Plesset equation. The extent of these lung dynamics was dependent on physical confinement, which in real underwater blast conditions is influenced by factors such as rib cage properties and donned equipment. Findings demonstrate a potential causal mechanism for implosion injuries, which has significant implications for the understanding of primary blast lung injury due to underwater blast exposures.

## Introduction

Since the early days of naval warfare in the 19th century, underwater explosions have been responsible for serious injury or even death (1, 2). Underwater explosive devices such as mines, torpedoes, and depth charges were increasingly common in the early 20th century (3). In World War II alone, over 1,500 casualties related to underwater blast were documented (4). While there have been fewer documented cases of injuries due to underwater blasts in recent decades (5), naval warfare has become one of the major emerging battlespaces of the future (6). As a result, injury or death to service members exposed to underwater blasts has the potential to be more prevalent in the future.

While extensive work has been undertaken to investigate safe and lethal exposure levels under primary blast exposure in air (7–10), the translation of these limits to underwater blast injury risk is challenging due to fundamental differences in shock wave characteristics between water and air. Shock waves propagate farther and faster in water due to the higher density of water and the corresponding increased speed of sound. When these shock waves reflect off the water surface or ocean floor, they can produce either constructive or destructive interference (11). Furthermore, the underwater detonation can cause gas bubble cavitation, which generates additional shock waves (11). Due to the similar densities between humans and water, underwater shock waves are able to transmit more energy to the organs, posing a greater risk to humans (12, 13). Conversely, the human body reflects more energy from shock waves in air (12).

Gas-containing organs are most vulnerable to underwater primary blast injury (14) due to the sudden decrease in material density and sound speed at the gas-tissue interface, which results in increased shock wave energy deposition (12). Trauma to the lungs is particularly lethal (15–17), as it can result in pulmonary hemorrhage and contusions, gas embolisms, and pneumothorax, among others conditions (16, 18, 19). Mechanisms of underwater blasts injuries are thought to closely resemble those suggested for air blasts (20), i.e., spallation, implosion, and inertia (16, 21, 22). However, these mechanisms are poorly understood due to the lack of experimental evidence.

Historically, underwater blast injury studies have sought to establish exposure guidelines for the lungs (13, 20). Data that inform these guidelines are based on air blast, unscaled animal models, computational models, medical case reports, clinical experience, or even, speculation (13, 20). The broad range of methods has led to highly inconsistent guidelines, without a consensus exposure metric (e.g., peak pressure or impulse, or charge weight and range). Most importantly, these guidelines are not founded on well-characterized experimental data for humans, which is critical for the establishment of relevant and precise injury guidelines. Until a robust mapping between underwater explosions and human injury is established, military missions will continue to expose service members to underwater blast with an unknown risk of injury or death.

To address the critical need for high-fidelity human lung data, a series of shock tube experiments were conducted where isolated human lungs were exposed to underwater shock waves in a water chamber. The pressure and volumetric response were measured with a combination of pressure sensors and high-speed video, and compared to equivalent measurements from solid organs, i.e., the liver and spleen. Finally, an analytical model based on the Rayleigh-Plesset equation was utilized to further explain the mechanisms that lead to the observed pressure-volume changes and to inform future injury risk metrics.

## Materials and methods

### Shock Tube Setup

A shock tube, designed to simulate blast loading pressure profiles, generated short duration underwater overpressure waves to expose submerged organs (Fig 1). The shock tube was divided into four sections; the driver, driven, diffuser, and water chamber (Fig 1A). The driver and driven section are separated by a diaphragm that prevents the flow of pressurized helium from the driver to the driven section. The diaphragm ruptures once a threshold pressure is reached, which produces a shock wave as the pressure wave travels along the driven section. Pressures were measured by a PCB sensor (PCB113B21, PCB Piezotronics, Depew NY). The diaphgram condition was calibrated to deliver repeatable rupture pressures. The pressure wave is then radially expanded from 0.15 m to 0.43 m in diameter by an air-filled conical diffuser as it travels towards the water filled chamber. The diffuser and water-filled chamber are separated by a rubber diaphragm that ensures that water does not enter into the driven section, but still allows for transmission of the shock wave into the water chamber. Tests were run at two burst pressures of 350 kPa and 700 kPa for the liver and the spleen, and 350 kPa and 525 kPa for the lung. There was a discrepancy of the higher burst pressure between the liver and spleen, and lung due to manufacturing and storage differences of the diaphragm materials. Three repeat tests were conducted run at each burst pressure for each specimen.

**Fig. 1.**
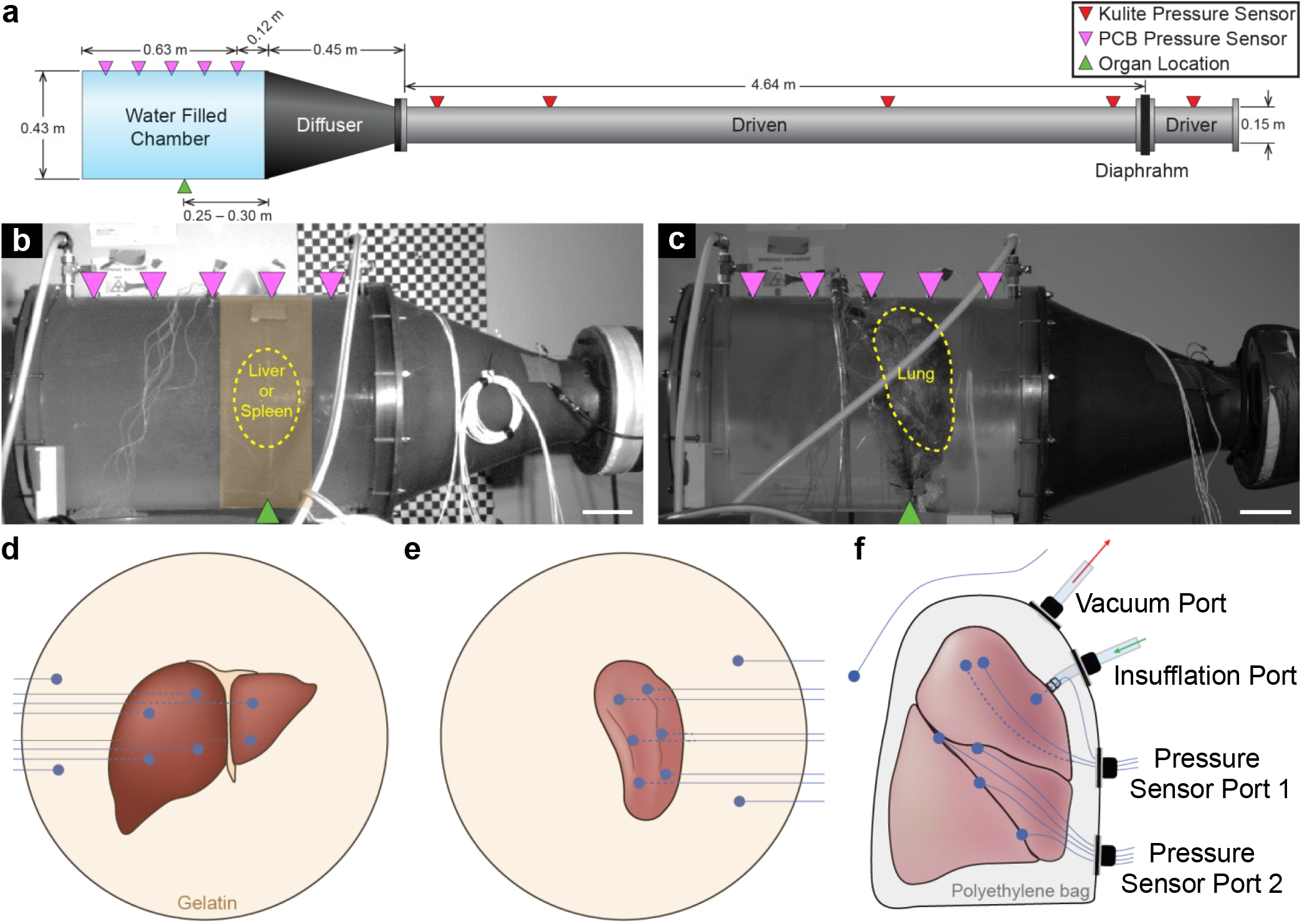
Experimental setup and organ preparation method for applying overpressure to organs within a water-filled chamber. (A) Schematic illustrating the key components of the shock tube, including pressure sensors (red downward-point triangles) for measuring the pressure evolution of the shock wave in the driver and driven section, and pressure sensors (magenta downward-pointing triangles) for measuring the overpressure applied to the organ in the water chamber. Lateral images shows the lateral placement (green triangle) and outline (yellow dotted line) of the (B) liver and spleen, and (C) lung within the water chamber. Scale bar, 0.1 m. To prepare the organs for testing, the (D) liver and (E) spleen are encapsulated in a gelatin puck and instrumented with six pressure sensors (blue circle) inserted into the parenchyma and two reference pressure sensors inserted into the gelatin. (F) The right and left lungs were encapsulated in a polyethylene bag installed with a lung insufflation port, a suction port for removing air leakage from the lung, and two sensor ports that pass-through eight pressure sensors (blue circle), with six located under the visceral pleura, one inserted into the main bronchus, and one positioned in the water chamber as a reference measurement.

The burst pressure was measured by a Kulite pressure sensor (HKS-37, Kulite Semiconductors, Leonia, NJ), installed through the wall of the driver section. The dynamics of shock wave pressure as it propagates along the driven section was measured by an additional five Kulite pressure sensors placed at predetermined intervals. The pressure in the water chamber was measured by five piezoelectric PCB pressure sensors (PCB113B21, PCB Piezotronics, Depew NY), installed through the wall of the water chamber. All pressure data was collected at 1 MHz using a 16-bit high speed data acquisition system (DEWE 801; Dewetron, Wakefield, RI). Pressure values are relative to atmospheric pressure. Lateral images of the dynamic events during shock wave propagation in the water chamber were recorded by a high-speed camera (v711, Phantom, Wayne, NJ) at 2,000 fps. The last test series for the lung had an additional rear facing high-speed camera, which was used to compute the dynamic volume change due to the shock wave.

### Specimen Preparation

Tests were performed on the liver (*N* = 4), spleen (*N* = 4), left lung (*N* = 2), and right lung (*N* = 2) of six human cadaveric specimens obtained through the Maryland State Anatomy Board with an age ranging from 61 to 78 years. Informed consent was obtained from the donor or next of kin. All donors were screened to avoid any medical issues that would affect the mechanical properties of the tissue, e.g., cancer and chronic obstructive pulmonary disease. Specimens were fresh-frozen at -20 °C and stored until testing. Prior to preparation, specimens were thawed for at least 8 hours at 4 °C. The solid organs, the liver and spleen, were encapsulated in a gelatin puck (Fig 1D,E). Gelatin was chosen to correspond with the shock impedance properties of soft tissue (23–25) and water (26). A 10% w/v gelatin puck was created from 250 Bloom A Gelatin powder (Knox, Sioux City, IA) according to the protocol outlined by Fackler and Malinowski (27). Gelatin powder was rigorously mixed into cold water at 7 – 10 °C and subsequently heated until the gelatin was completely dissolved. The gelatin solution, totaling 12 L, was then poured into a cylindrical mold with the same diameter as the water chamber and allowed to cure for at least 8 hours at 4 °C . Subsequently, the organ was placed into a cavity in the shape of the organ, which was cut from the surface of the gelatin. An additional 12 L of gelatin solution was poured into the mold to encapsulate the rest of the organ. The gelatin supported the organs to approximate physiological geometry for the duration of the experiment. The final dimensions of the gelatin puck were 0.41 m in depth and 0.43 m in diameter. To measure the dynamic pressure due to the pressure wave in the water chamber, six fiber optic pressure (FOP) sensors (FOP-M-PK, FISO, Quebec, QC, Canada) were inserted into the parenchyma of the liver and spleen through a hollow insertion tube. Two additional FOP sensors were placed into the gelatin to serve as references for computing the incident pressure. The location of the FOP sensors for the liver and spleen are shown in Fig 1D,E. A similar procedure was initially repeated for the lungs. However, air leakage during potting and testing compromised the mechanical integrity of the gelatin. Additionally, it was not possible to confirm the insufflation of the lungs during testing since the cured gelatin is opaque.

To overcome these issues with air leakage, a novel encapsulation method for underwater testing of the lungs was developed. The lung was inserted into a polyethylene bag with four liquid-proof ports installed. Two ports served for insufflation and vacuum, and two ports served to insert sensors (Fig 1F). The lung was insufflated during testing by a pump that delivered air at pressures ranging from 5 – 10 kPa through a vinyl tube that passed through the insufflation port and connected to a barbed polyethylene fitting sutured to the bronchus. These insufflation pressure ranges were based on typical mechanical ventilation pressures (28). The vacuum port was attached to a pump with a vinyl tube and evacuated the air leaking out of the lungs during testing. To measure the dynamic pressure response of the lung, seven FOP sensors were inserted under the visceral pleura, and one FOP sensor was inserted into the main bronchus. An additional reference pressure sensor was placed next to the lung in the water chamber. This encapsulation method provided three key advantages: 1) precise control of lung insufflation, 2) removal of air leaking from the lung to the water chamber, and 3) full visibility of the lung, allowing the capture of highspeed video.

Prior to testing, organs were placed into the chamber with no water and positioned along the radial center of the chamber and approximately 0.25 – 0.30 m from the end of the diffuser. To position the lungs for testing, a thin plastic net was anchored to the chamber walls. The sensor cables and tubing for the lung were passed through water-tight ports at the top of the water chamber and the chamber was subsequently filled with water until no air was present. Photos of the nominal pretest position of the liver, spleen, and lungs are shown in Fig 1B,C.

### Data Processing and Analysis

#### Pressure Measurements

Data was processed and analyzed using MATLAB 2022b (Mathworks, Natick, MA). All pressure measurements except those made by the reference pressure sensor next to the organ were filtered with a zero-phase 4-pole Butterworth low-pass filter with a cutoff frequency of 50 kHz. Peak organ pressure was determined by computing the local maxima within 30 ms of the trigger and subsequently verified through visual inspection of the data trace. The dominant frequency of the pressure response of the organ was computed by averaging Welch’s power spectral density (29) across all pressure sensors and subsequently selecting the frequency with the highest power.

To quantify the pressure dose to the organ, incident pressure was computed by subtracting the reference pressure measurement from the chamber pressure measurement closest to the diaphragm. Prior to subtraction, the reference pressure measurement was low-pass filtered with a zero-phase 4-pole Butterworth with a cutoff frequency of 1 kHz to preserve the transient response of the incident pressure wave. This method of computing the incident pressure was chosen in order to overcome limitations associated with pressure measurements collected in an enclosed rigid chamber, where the pressure response of the organ alters the measured chamber pressure due to the relative incompressibility of the water. The subtraction procedure isolates the transient pressure dose from the organ pressure response. S1 Fig shows an example of pressure waveforms of the incident pressure computed using this method.

A two-sided hypothesis test was performed to examine the linear association between the peak pressure and incident pressure, with a significance level of *p* < 0.05 based on the computed t-statistic of the slope term.

A upphapiro-Wilk test was performed to assess the normality of both the peak pressure range and the dominant frequency. To identify significant pairwise differences across organs, a Kruskal-Wallis test followed by a post-hoc Dunn-Sidak test was conducted, with a significance level set at *p* < 0.05.

#### Lung Volume Measurements

The volume of the lung (*V*) for the last test series was computed from the high-speed video by approximating the lung for a single test as an ellipsoid. The equation for the lung volume and corresponding volumetric strain *ε*_*V*_ relative to the initial volume *V*_0_ is given by

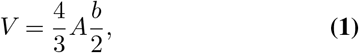

and

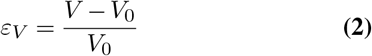

respectively, where *A* is the cross-sectional area of the lung computed by manually tracing the lung boundary from the lateral high-speed video and *b* is the distance corresponding to the minor axis of an ellipse that was manually fitted to the lung boundary from the rear high-speed video. Manual tracing was repeated every 3 ms for a total of 99 ms post diaphragm burst. The other test series for the lungs were not included in the volumetric analysis since they did not include high speed video of the chamber from the rear view. The volumetric strain rate was computed with forward finite difference.

#### Analytical Modeling of the Lung Dynamics

A modified version of the Rayleigh-Plesset (RP) equation (30, 31) was applied to understand the dynamic response of the lungs due to a transient pressure wave (32, 33). For this model, the lungs are assumed as a spherical gas bubble suspended in a spherical domain of incompressible liquid, which is confined by an elastic spherical shell. The choice of the gas bubble confinement was made to get a preliminary understanding of the confinement effects of the rib cage in humans. The following assumptions are made: (1) the shell inertia are negligible; (2) the gas bubble behavior follows a polytropic process and its pressure is uniform; (3) there is zero mass transport across the bubble interface; and (4) the dynamic viscosity and surface tension effects are negligible due the large dimensions of the bubble (i.e., *>* 10^−1^ *m*) (34). The equation of motion for the bubble is given by

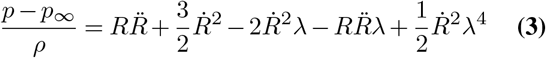

where *ρ* is the liquid density, *p* is the pressure inside the bubble, *p*_∞_ is the pressure of the liquid, *R* is the bubble radius with time derivatives 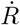 and 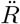,and *λ* is a dimensionless parameter defined as the ratio of *R* to the radius of the spherical shell *R*_S_ (i.e., *λ* = *R/R*_S_). The last three terms that contain *λ* are the modification to the classic RP equation, which can be obtained by setting *R*_S_ = *∞*. Due to the liquid incompressibility, *R*_S_ is related to *R* via volume conservation by

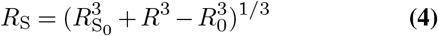

where 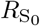 and *R*_0_ are the initial shell radius and bubble radius, respectively. While other versions of the RP equation (35, 36) that can better generalize to other boundary conditions, this version of the confined RP was chosen due to the inclusion of key parameters without undue complexity. Other models of the lungs that account for structure, material properties, and geometry (36–39) were considered, but complexity beyond the needs of this study placed these models out of scope.

Eq (3) was numerically solved with MATLAB “ode45” for *p* and *R* as a function of time, *t*, given an incident pressure *p*_*∞*_, 4 *b* defined as an instantaneous pressure increase with amplitude *p*_A_ from initial pressure *p*_0_ with duration *τ*, i.e.,

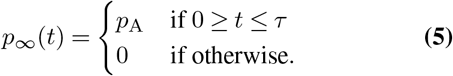

The volumetric strain was computed with Eq (2), but assuming the volume of a sphere, i.e., *V* = 4*/*3*πR*^3^.

The parameters for the model were chosen to best match the experimental data. The initial gas bubble radius *R*_0_ was set to 0.092 m based on the average effective radius of lung prior to shock wave arrival as computed from the volume estimated with equation (1). The effective polytropic index for the lung was set to 1 based on Wodicka et al. (40). The liquid density *ρ* was set to that of water, i.e., 997 kg/m^3^. Similar to the experimental data, all pressures are relative to atmospheric pressure.

## Results

### Organ pressure response waveform

A representative pressure response measured by sensors embedded throughout the organ in various locations, along with the corresponding incident pressure waveform is shown in Fig 2. The average peak incident pressure was 68 kPa, 86 kPa, and 113 kPa resulting in a peak organ pressure of 88 kPa, 106 kPa, and 119 kPa for the lung, liver, and spleen, respectively. The pressure response of the lung shows large regional differences in the pressure magnitude (Fig 2A). In contrast, the pressure response of the liver and spleen (Fig 2C,D) are tightly grouped, indicating minimal regional differences in pressure magnitude. The oscillatory behavior was markedly higher for the liver and spleen compared to the lung. The morphology of lung pressure was markedly different, in which the positive pressure peaks were shorter and greater in magnitude compared to the longer negative pressure troughs. The insets in Fig 2 provide a more detailed version of the pressure waveforms, and reveals that the liver and spleen exhibit a considerably fast, approximately 2 ms, pressure rise time, compared to the lung with a rise time of approximately 10 ms. The high-frequency and transient behavior of the incident pressure wave is not present in all of the organ.

**Fig. 2.**
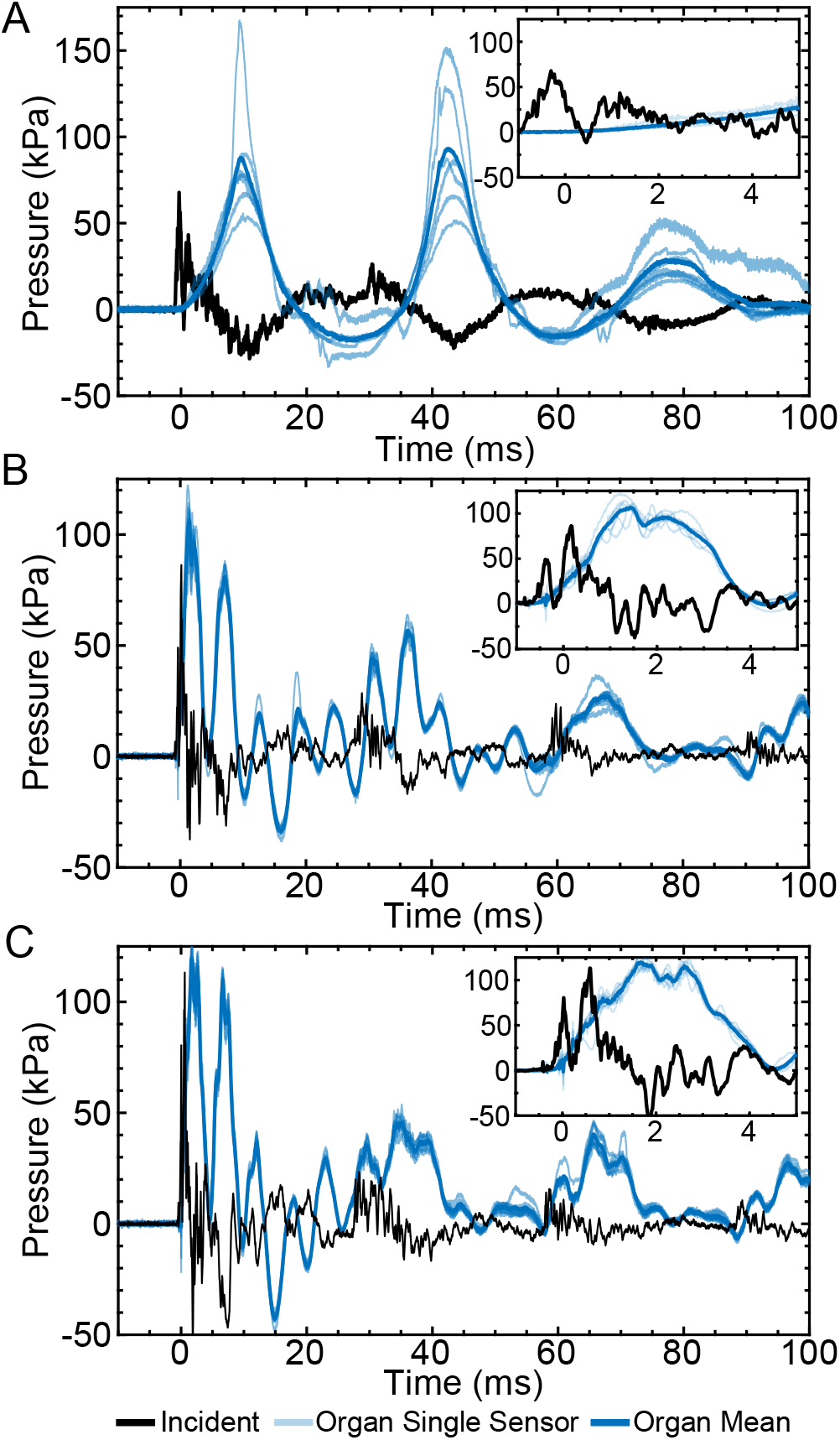
Representative organ pressure response waveforms. (A) lung, (B) liver, and (C) spleen pressure response due to incident pressure (black line), The light blue line shows the individual organ sensor measurements, while the blue line represents the temporally averaged response. Inset shows the first 5 ms of the pressure responses.

### Features of the organ pressure response

The relationship between peak incident pressure, and the mean and maximum peak organ pressure is shown in Fig 3A,B. The mean peak lung pressures due to incident peak pressures of 53 kPa– 108 kPa ranged between 55 kPa – 147 kPa, but was not significantly correlated to peak incident pressure (*R*^2^ = 0.16, *p* = 0.06). Conversely, the liver and spleen exhibited a wider range of mean peak organ pressures from 79 kPa to 190 kPa, due to greater burst pressures, which produced a greater down-stream incident pressures from 46 kPa to 177 kPa compared to the lung. A significant positive correlation was observed between peak incident pressure and peak organ pressure for the liver (*R*^2^ = 0.81, *p* < 0.001) and spleen (*R*^2^ = 0.75, *p* < 0.001). Maximum peak lung pressures were considerably higher than mean peak lung pressures, and ranged from 68 kPa to 394 kPa, but no significant correlations were observed (*R*^2^ = 0.16, *p* = 0.06). The maximum peak pressure of the liver and the spleen were similar to mean peak pressure and ranged from 83 kPa to 252 kPa. Similar trends across organs were observed with incident impulse, which ranged from 28 to 196 N*·*ms, likely due to correlation between the peak incident pressure and the associated impulse (20). Maximum peak organ pressure significantly correlated with the peak incident pressure for the liver (*R*^2^ = 0.80, *p* < 0.001) and spleen (*R*^2^ = 0.75, *p* < 0.001). These differences between mean and maximum peak organ pressure were also be observed by computing the regional range of the peak organ pressure on a per test basis (Fig 3C). The lung exhibited a significantly higher peak organ pressure range (median = 103 kPa) than either the liver (median = 19 kPa, *p* < 0.001) or the spleen (median = 22 kPa, *p* < 0.001) (Fig 3C). No significant differences were observed between pressure ranges for the liver and spleen (*p* = 0.82). The lung exhibited a significantly lower dominant frequency response (median = 28 Hz) compared to the liver (median = 176 Hz, *p* < 0.001) and the spleen (median = 198 Hz, *p* < 0.001) (Fig 3D). The dominant frequency between the liver and the spleen were significantly different (*p* < 0.001).

**Fig. 3.**
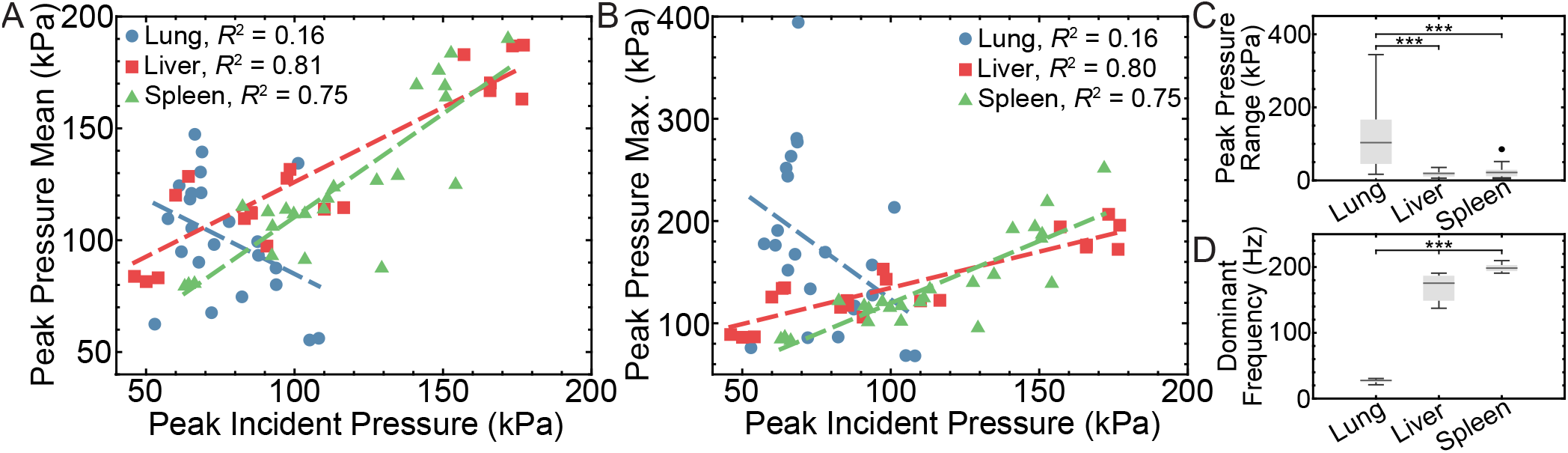
Organ-specific pressure response characterization. (A) Mean and (B) maximum peak pressure response of lung (blue circle), liver (red square), and spleen (green triangle) with corresponding line of best fit (blue, red, and green dashed lines, respectively) as a function of peak incident pressure. Box-and whiskers plots of the (C) peak pressure range and (D) dominant frequency of the pressure response for liver (*n* = 20, *N* = 4), spleen (*n* = 24, *N* = 4), and lungs (*n* = 23, *N* = 4) showing the median (black line), mean (plus), and interquartile range (gray box). Outliers (circle) are 1.5 times the interquartile range either above or below the non-outlier maximum or minimum shown as whiskers. \*\*\**p* < 0.001.

### Volumetric response of the lung

The lung underwent large volumetric strains and strain rates due to the pressure wave in the water-filled chamber compared to the liver and spleen (Fig 4). The minimum and maximum volumetric strain for the specimen shown in Fig 2A was -24.0% and 15.6%, respectively (Fig 4A). The maximum and minimum volumetric strain rate was 43.3 s^−1^ and -41.7 s^−1^, respectively (Fig 4B). The volumetric strain oscillations occurred at the same dominant frequency as the pressure oscillations shown in Fig 2A. The corresponding lateral high-speed images of the lung in the undeformed, most compressed, and most expanded state of the lung are shown in Fig 4C,D. In the most compressed state, the lung surface deformed nonuniformly.

**Fig. 4.**
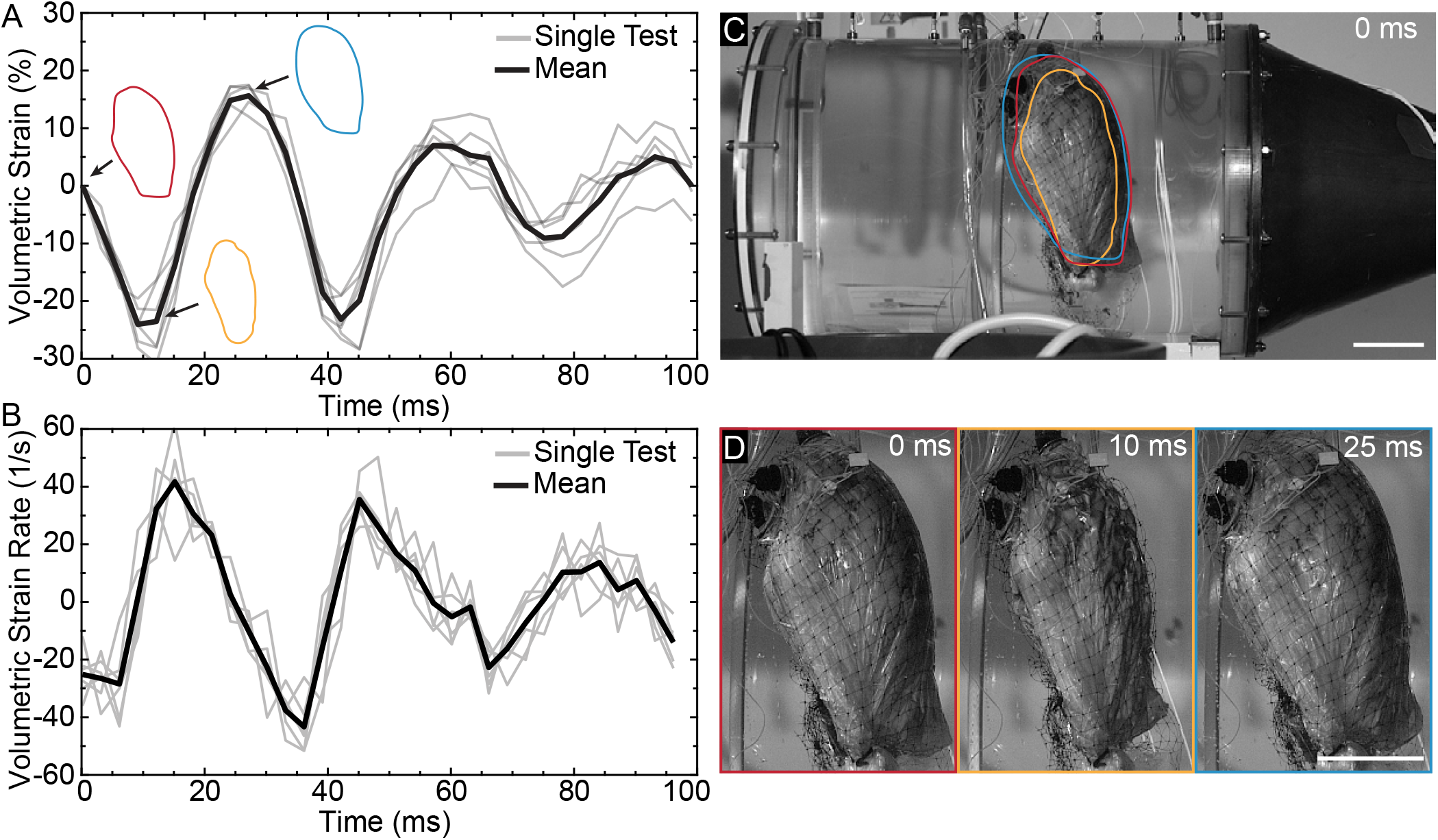
Volumetric deformation of the lung due to shockwave exposure. (A) Volumetric strain and (B) strain rate time series of the lung with six tests (gray) and corresponding mean (black). Outlines of the lung are shown in the undeformed at 0 ms (red), most compressed at 10 ms (yellow), and most expanded at 25 ms (blue) state. (C) Lateral image in the undeformed state with an overlay of the lung outlines shown in (A). (D) Enlarged images of the lung shown in (C). Scale bar, 0.1 m.

### Analytical model of the lung pressure-volume response

A confined Rayleigh-Plesset (RP) equation was solved to understand the driving force behind the pressure-volume response of the lung due to a transient pressure pulse. For this model, the lungs are assumed to be a spherical gas bubble with initial radius *R*_0_ suspended in a spherical domain of incompressible liquid confined by a spherical shell with radius *R*_S_. Fig 5 shows solutions for the bubble pressure (*p*) and volumetric strain *ε*_V_ for a range of different incident pressures amplitudes (*p*_A_) and durations (*τ*), and for different bubble confinement (denoted as the ratio of *R*_S_ to *R*_0_). The waveform morphology of bubble pressure exhibited shorter duration positive pressure peaks with larger magnitudes compared to the longer negative pressure troughs, which were more pronounced with higher incident pressures (Fig 5A). The corresponding volumetric strain of the bubble was inversely related to the bubble pressure due to the gas behavior following a polytropic process. The maximum bubble pressure and volumetric strain scaled nonlinearly with both the incident pressure amplitude and duration (Fig 5B,C) and impulse (S2 Fig). At incident pressure durations of 0.1 ms and 1 ms, the maximum bubble pressure was less then incident pressure amplitude. However, at higher incident pressure durations of 10 ms, the maximum bubble pressure exceeded the incident pressure amplitude by 2.1 to 12.6 times. The critical incident pressure impulse that produces greater bubble pressures than incident pressures is approximately 270 kPa ms. As the bubble becomes more confined (i.e., *R*_S_*/R*_0_ → 1.1), the maximum bubble pressures and volumetric strains increased by 3.3 to 15.9 times and 3.2 to 4.5 times, respectively (Fig 5D).

**Fig. 5.**
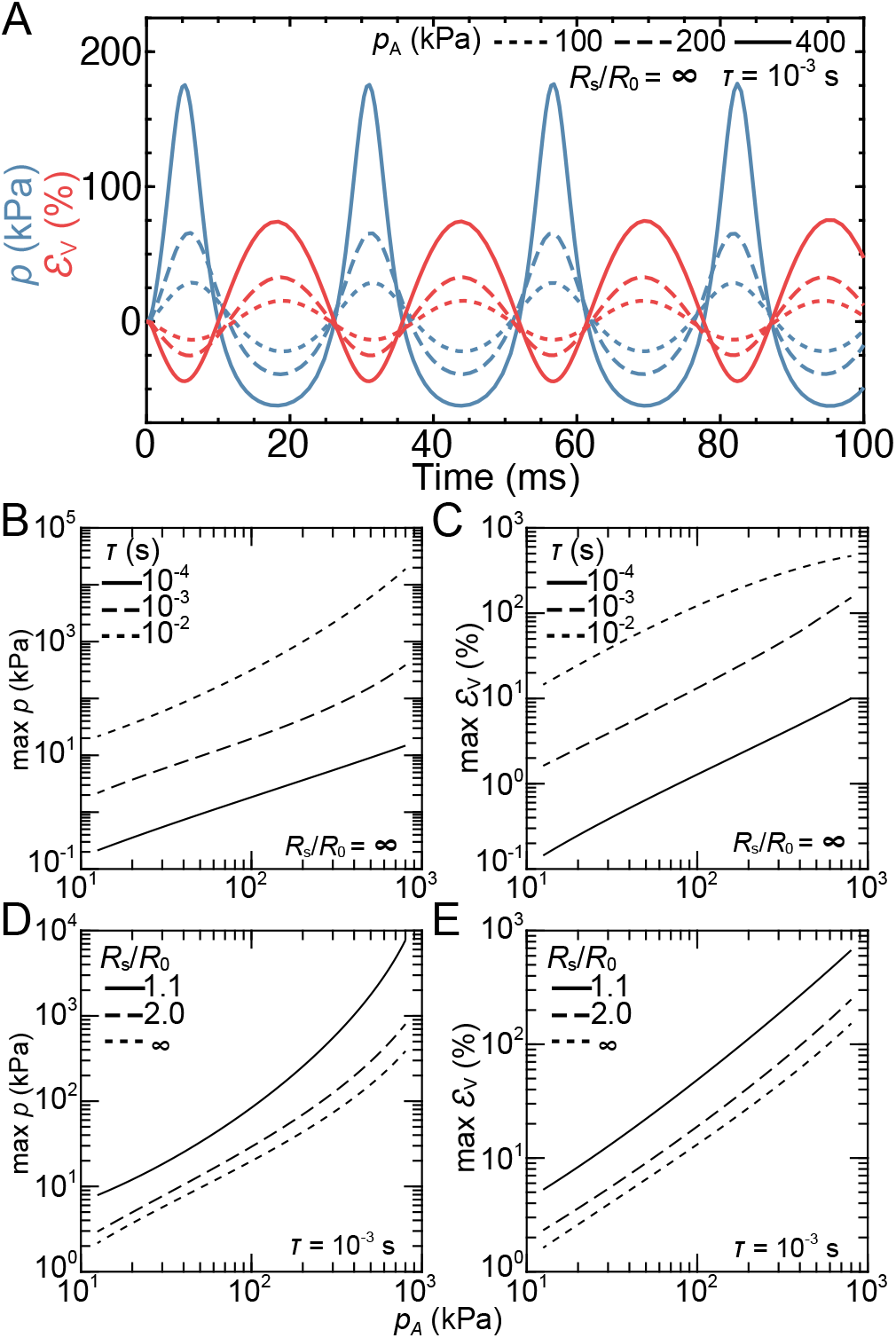
Analytical solution of the modified Rayleigh-Plesset equation of a spherical gas bubble with initial radius *R*0 within a spherical water chamber of radius *R*_S_. (A) Bubble pressure, (*p*; blue) and volumetric strain (*ε*_V_; red) waveforms due to a square pressure pulse with amplitudes *p*_A_ = 100 kPa (dotted), 200 kPa (dashed), and 400 kPa (solid) and duration *τ* = 10^*−*3^ s within an infinitely large water chamber, i.e., *R*_S_*/R*_0_ = . Maximum (B) *p* and (C) *ε*_V_ for increasing values of *p*_A_ and *τ* = 10^*−*4^ s (solid), 10^*−*3^ s (dashed), and 10^*−*2^ s (dotted) at *R*_S_*/R*_0_ = *∞*. Maximum (D) *p* and (E) *ε*_V_ for increasing values of *p*_A_ and *R*_S_*/R*_0_ = 1.1 (solid), 2.0 (dashed), and *∞* (dotted) at *τ* = 10^*−*3^ s.

## Discussion

While there been many attempts to establish injury guidelines for the human lungs exposed to underwater blast (13, 20), the criteria remain highly variable due to a lack of sufficient human data to reveal the underlying injury mechanisms. To address this gap, a series of shock tube experiments that subjected isolated lungs to shock waves in a water chamber were conducted. Experiments were repeated with the liver and the spleen to compare lung response to those of solid organs. Lastly, this study utilized an analytical model based on the Rayleigh-Plesset (RP) equation to isolate the effect of air on lung response and to understand the mechanisms of lung deformation.

Upon analyzing the pressure measurements (Fig 2A), transient spikes in lung pressure were not observed with a dunitudes with longer negative pressure troughs (Fig 2A) repeating at approximately 28 Hz (Fig 2D). Unlike the solid organs, the peak pressures associated with these cycles exhibited large test-to-test variations that did not correlate with peak incident pressures (Fig 3A,B). In some tests, the measured peak pressure greatly exceeded peak incident pressure. This finding provides further evidence that the pressure response is not dominated by the shock wave front.

The pressure cycles in Fig 2A and inversely associated volumetric strains (Fig 4B) are indicative of the thermodynamic processes of gases (43). To gain insights into this interesting PV behavior, we solved a RP equation where a spherical gas bubble within a domain of incompressible liquid subject to a short-duration pressure square wave (30, 31). Fetherston et al. solved a similar equation to understand the dynamics of marine mammal lungs when exposed to underwater blast (36). Although this model oversimplifies the complexities of lung composition, material properties, and structure, the PV time series (Fig 5A) exhibits waveform morphologies that are remarkably similar to those of the lung (Fig 2A and Fig 4B). These morphological similarities provide evidence that the bulk PV response of the lung is due to the compression of the contained gas, which is initiated by the shock wave. One possible mechanism for how the shock wave initiates lung compression is that the external water-tissue interface has a small acoustic impedance mismatch, so the reflection from the water-tissue interface is small, allowing more energy to be transmitted into the body. However, at the interface between the pleural cavity and the lung, the acoustic impedance ration similar to the incident pressure waveform, suggesting that shock wave front propagation through the lungs is severely attenuated. This attenuation is likely due to the unique structure of the lung, which is composed of many microscopic air sacs. Each air sac acts as a high acoustic impedance solid-gas interface that diffracts and reflects the shock wave front. At a macro-scale, these events superimpose to severely and quickly dissipate the energy of the shock wave front. This proposed dissipation mechanism is similar to the well-characterized shock wave attenuation mechanisms in foams (41, 42). Unlike the lungs, the solid organs exhibit a more transient pressure response that lasts approximately 4 ms (Fig 2B,C), further highlighting the influence that the structure and composition of the organ have on attenuating the shock wave front.

Despite substantial shock wave attenuation, the lungs still underwent large pressure cycles characterized by larger magnitudes with shorter positive pressure peaks, and smaller mag-mismatch is large, leading to substantial energy deposition at the lung surface, which then initiates a bulk PV response. This proposed mechanism of lung compression in underwater blast exposure is substantially different from the mechanism of lung compression in air blast exposure as modeled by Stuhmiller (44), due to the difference in the surrounding fluid. In Stuhmiller’s analysis, the air-to-tissue interface reflects the blast wave, resulting in momentum transfer to the outer tissues of the chest and abdomen. Resulting motion of the chest wall and diaphragm are then used to develop a model for lung compression. Another possible mechanism for how the shock wave initiates lung compression can be observed in studies involving foams, where heavily attenuated shock waves convert to high-pressure compression waves causing foam compaction (45). For both initiation mechanisms, we expect that these PV cycles are also present when the lung is exposed to air blast, but with smaller amplitudes due to weaker acoustic coupling between the torso and the air compared to coupling with water (12, 13), and higher frequencies due to air having less inertia than the surrounding water.

Confinement on the lungs by the rib cage plays a critical role in PV response. To understand these effects, a solution to the modified version of the RP equation that accounts for confinement was solved by enclosing the gas bubble and surrounding liquid with an elastic shell (32, 33). By accounting for confinement, bubble pressures substantially increased by approximately 10 to 15 times when the bubble was in a shell that is 10% larger than its original radius (Fig 5D). Although we expect the corresponding volumetric strain in real scenarios to decrease with confinement in humans, our model shows the opposite (Fig 5E). This discrepancy is attributed to the treatment of the elastic shell in Eq (4) as variations of bubble volume were accommodated by modifying the shell radius. From an injury perspective, a decreased volumetric strain is desirable. Yet, this accommodation comes at the cost of inducing higher alveolar pressures, which could lead to increased forced air emboli into the capillary (46). These increased pressures could also lead to local tissue shearing when the soft lungs impinge on the stiffer rib cage, which is consistent with clinical observations of rib markings on the lungs following blast injury (47). The effects of lung confinement are likely to vary based on the individuals rib cage stifness and geometry, as well as donned personal protective equipment, or occupation specific equipment, which may further restrict the lungs.

These findings have significant implications for our understanding of the injury mechanisms for lungs and other gas-containing organs exposed to underwater blast. While it is currently believed that the mechanisms of lung injury in underwater blasts closely follow those of air blasts (20), i.e., spallation, implosion, and inertia (21), the extent of damage caused by these mechanisms remains unknown despite numerous studies on air blast injuries (16). Among these mechanisms, implosion forces are the most consistent with the observed lung response in this study, resulting in rapid compression and expansion of gaseous content. At the alveolar length scale, compression can cause the alveolus to collapse and result in atelectasis (48), while pneumothorax can occur at the length scale of the lung (18, 19). Rapid lung expansion can cause alveolar and capillary overstretching and rupture, or the driving of extravascular fluid into the alveolar space, causing pulmonary oedema and hemorrhage (16). These injuries may not present uniformly throughout the lung based on regional pressure differences (Fig 2 and Fig 3C) that are due to the heterogeneous structure of the lung. Previously, the implosion mechanism was first postulated by Forbes in 1812 (49), later described by Schardin in 1950 (21), and conceptually modeled by Ho in 2002 (46). Yet, to the best of our knowledge, this study is the first to present experimental evidence of this mechanism, and with direct visualization of lung volume over the course of events.

Peak incident pressures and associated impulses measured in this study fall within the reported range of previous studies (13, 20). However, it is difficult to determine the severity of injury that would be obtained in this study with any granularity based on the large spread in the injury criteria (13, 20). This variability in reported data is likely due to the variety of approaches that have been used to develop these criteria, each with its own significant limitations (13). The peak incident pressures and impulses measured in this study are most likely above safe levels based on an animal study conducted by Richmond et al. (50, 51), but below 50% lethality based on a study by Lance et al. (20) that combined field injury data with computational predictions of incident pressures and impulses. It is important to note that the injury criteria developed in these studies are based on incident pressure and not the lung pressure, which can reach up to approximately six times the peak incident pressure (Fig 3B). These internal pressures should be an important factor in the development of future injury criteria, as they are a more accurate representation of tissue level loading that directly leads to injury.

## Conclusion

This study provides the first directly observable experimental data and characterization of human lung dynamics when exposed to underwater blast. We found that the shock wave front was severely attenuated by the high acoustic impedance gas-solid microstructure of the lung, similar to gas-filled foams (41, 42). However, the shock wave front initiated large bulk PV cycles that are distinct from the solid organs. By solving the RP equation, we show that these large PV cycles are due to the compression of contained gas, which follows a classic thermodynamic process (43). By further modifying the RP equation to include physical confinement, we find that the PV cycles are also highly depending on physical confinement, which is dependent on the rib cage properties and may be modified by donned equipment. These findings have significant implications for our understanding of the proposed injury mechanisms both for underwater and air blast exposures, in that they provides the first direct evidence of the implosion injury mechanism, which has was first proposed in 1817 by Forbes (49) and has been expanded on over the course of over two centuries (21, 46).

A number of future studies are needed to fully characterize lung dynamics during blast and their role in injury. In this study, isolated lungs were placed in a chamber that is not fully representative of human blast exposure in an open body of water. Future studies should characterize the dynamics of the lungs with a combination of experimental models. These studies should include postmortem human subject experiments to better understand the effects of the ribcage, and animal experiments to better characterize injury *in vivo*. To fully understand the injury mechanisms on the alveolar length scale, more detailed *in vitro* and *in vivo* models are needed in conjunction with higher resolution imaging techniques (52). Future studies should aim to create underwater shock wave loading in larger, open water scenarios where exposure occurs at depth, and near the surface to understand effects of shock wave rarefaction (13, 20). These shock waves should be generated with underwater explosives to better represent real world exposure to blast, and should cover a larger range of incident pressures to form a basis of comparison with previous injury criteria (20).

Higher fidelity computational models of the lungs exposed to underwater blast are critical to understanding the injury mechanisms and designing protective measures. Our study involved the use of the RP equation to create an analytical model of the lung. However, this equation oversimplified the composition, material properties, and structure of real lungs, resulting in PV responses that were different from the test data. Peak pressures and volumetric strains, as well as their rates of decay, are different than the test data. Specifically, by the third PV cycle, the maximum PV of that cycle has decreased by over 50% (Fig 4B). We believe that these discrepancies are based on the need to explicitly include sources of energy loss. For example, this model does not account for dynamic viscosity of the liquid and bubble surface tension (31), but we believe that these factors are negligible due to the larger dimensions of the bubble (34). Additionally, the model does not account for the viscoelastic nature of the lung (53, 54), which would significantly affect both the peak PV and subsequent decay. Future studies should build on the history of high fidelity finite element models used for blast (39, 44, 55–59) to better understand the unique PV response. However, these models must be validated against high-fidelity human data collected in underwater blast scenarios similar to those presented in this study.

## ACKNOWLEDGEMENTS

The authors would like to thank Andrew Merkle and Robert Armiger for their management of the overarching project under contract # W81XWH-09-2-0168 and Catherine Carneal for manuscript review.

## AUTHOR CONTRIBUTIONS

E.B.-K, T.P.H. and C.K.D. designed the experiments. E.B.-K consolidated the experimental data, conducted the data processing and statistical analyses of the pressure data, developed the analytical lung model, and interpreted the results. J.M.D. processed the lung volumetric data. E.B.-K and A.S.I. conducted shock tube experiments. E.B.-K drafted the manuscript. J.M.D., K.A.O., and C.K.D. contributed to the preparation of the manuscript. All authors reviewed the manuscript.

## Supporting information

**S1 Fig.**
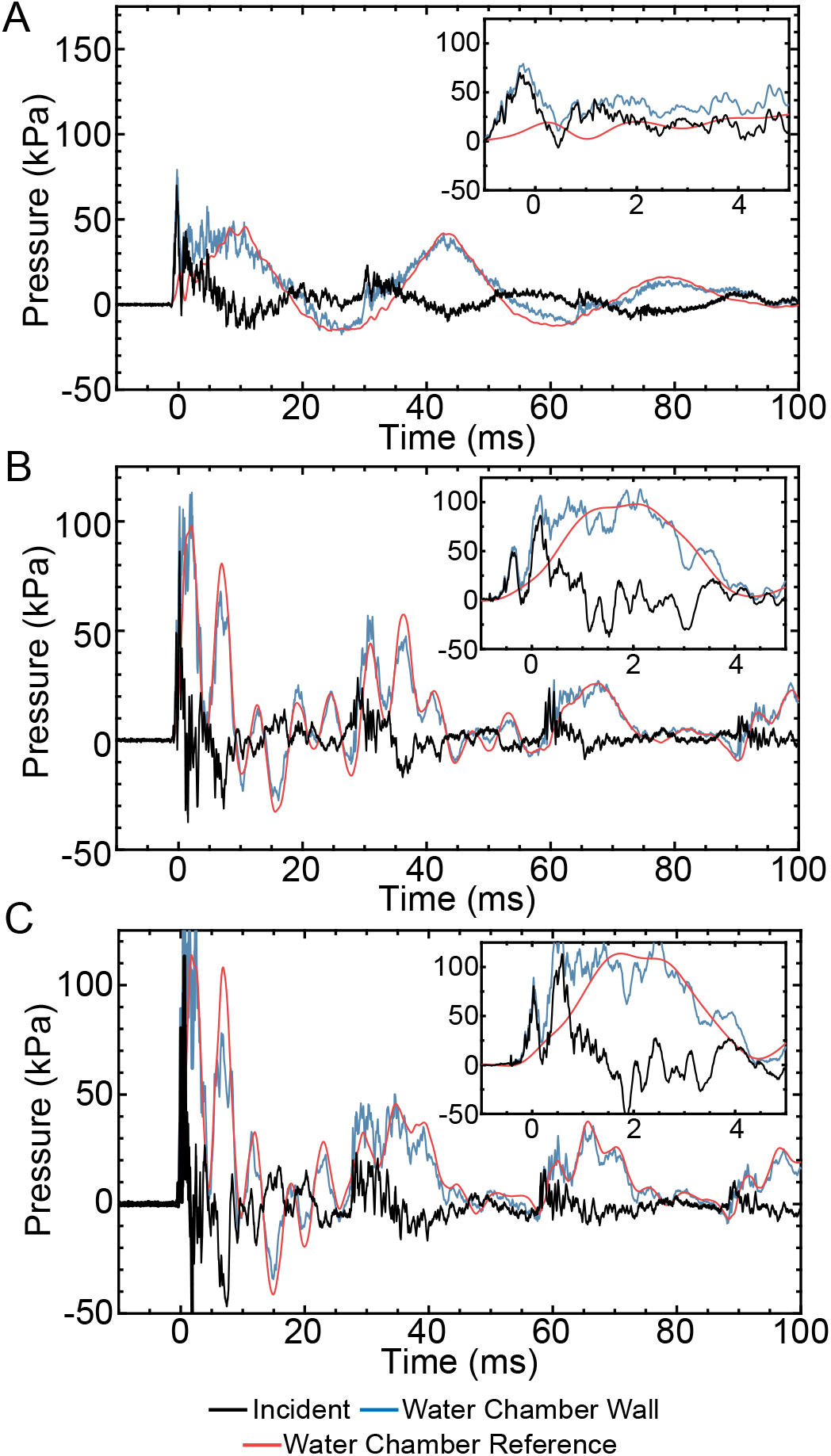
Representative pressure waveforms to illustrate the incident pressure calculation. The incident pressure (black) was computed by subtracting a filtered reference pressure measurement (red) from the pressure measurement made at the wall closest to the diaphram (green).

**S2 Fig.**
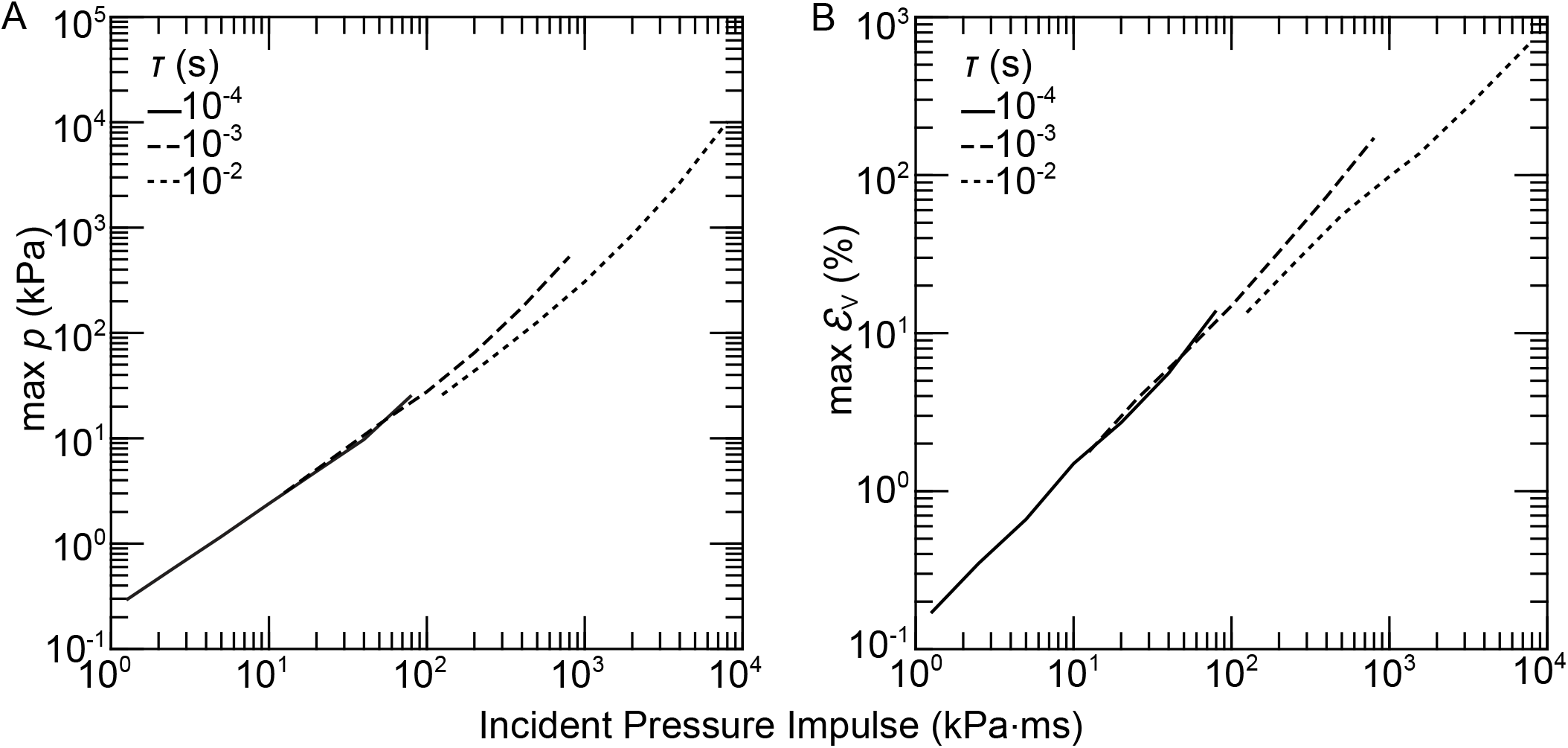
Bubble dynamics due to incident pressure impulse. Analytical solution of the Rayleigh-Plesset equation of a spherical gas bubble with initial radius *R*_0_ within a unconstrained spherical water chamber of radius *R*_S_ = *∞*. Maximum (A) bubble pressure *p* and (B) volumetric strain *ε*_V_ for increasing values of pressure impulse for incident pressure durations of *τ* = 10^*−*4^ s (solid), 10^*−*3^ s (dashed), and 10^*−*2^ s (dotted).

